# The ciliary gene *INPP5E* confers dorsal telencephalic identity to human cortical organoids by negatively regulating Sonic Hedgehog signalling

**DOI:** 10.1101/2021.06.06.447245

**Authors:** Leah Schembs, Ariane Willems, Kerstin Hasenpusch-Theil, James D. Cooper, Katie Whiting, Karen Burr, Sunniva M.K. Bøstrand, Bhuvaneish T. Selvaraj, Siddharthan Chandran, Thomas Theil

## Abstract

Defects in primary cilia, cellular antennas that controls multiple intracellular signalling pathways, underlie several neurodevelopmental disorders, but how cilia control essential steps in human brain formation remains elusive. Here, we show that cilia are present on the apical surface of radial glial cells in human foetal forebrain. Interfering with cilia signalling in human organoids by mutating the *INPP5E* gene leads to the formation of ventral telencephalic cell types instead of cortical progenitors and neurons. *INPP5E* mutant organoids also showed increased SHH signalling and cyclopamine treatment partially rescued this ventralisation. In addition, ciliary expression of SMO was increased and the integrity of the transition zone was compromised. Overall, these findings establish the importance of primary cilia for dorsal/ventral patterning in human corticogenesis, indicate a tissue specific role of *INPP5E* as a negative regulator of SHH signalling and have implications for the emerging roles of cilia in the pathogenesis of neurodevelopmental disorders.

## INTRODUCTION

The cerebral cortex is the largest and most complex part of the human brain and confers humans with their unique cognitive capabilities. The cellular and molecular mechanisms governing corticogenesis are under intensive investigation since cortical malformations underlie a number of neurodevelopmental disorders such as autism spectrum disorders (ASD) and intellectual disability (ID). The cortex develops from the most rostral part of the brain, the telencephalon. A patterning process subdivides the telencephalon early in development into distinct dorsal and ventral territories forming the cortex and the basal ganglia, respectively. The molecular basis for this process is well understood in mice (Rallu et al., 2002a) where dorsal progenitor cells express the Pax6 and Emx transcription factors and generate glutamatergic projection neurons either directly or indirectly via Tbr2+ basal progenitors (Englund et al., 2005; Götz et al., 1998; Warren et al., 1999). The ventral telencephalon is further subdivided into the caudal, medial and lateral ganglionic eminences (CGE, MGE and LGE, respectively) that express the Nkx2.1, Gsx2, Dlx2 and Olig2 transcription factors and produce a variety of different neuronal subtypes and oligodendrocyte precursor cells (OPCs). Notably, GABAergic cortical interneurons are formed in the MGE and CGE from where they migrate tangentially into the cortex (Marin and Muller, 2014). Generating these subdivisions of the telencephalon with their characteristic combinatorial expression of transcription factors is under the control of several signalling molecules (Theil et al., 2002). Members of the Wnt and Bmp gene families produced in the cortical hem promote cortical development whereas Shh signalling induces the formation of ventral telencephalic structures and cell types (Rallu et al., 2002a). Thus, a combination of signalling molecules and transcription factors direct murine corticogenesis, however, due to species-specific cellular features of the human brain (Florio and Huttner, 2014; Hodge et al., 2019) and due to a lack of suitable human model systems it remains largely unknown whether this mechanism is evolutionarily conserved between mice and humans. In particular, it has not been explored how telencephalic signalling is regulated at the cellular level during human cortex development and how defects in cell-signalling could contribute to neurodevelopmental disorders.

Primary cilia are small, microtubule-based protrusions from the cell surface that act as antennas in the detection and intracellular transduction of multiple cell-cell signals in development and tissue homeostasis. Defects in the function and/or structure of primary cilia underlie a group of syndromes commonly referred to as ciliopathies (Tobin and Beales, 2009). Ciliopathies are characterized by pleiotropic clinical features and many ciliopathy patients display severe neurological symptoms, most commonly ID (Valente et al., 2014). In turn, many candidate genes for ASD, schizophrenia, and ID affect primary cilia function (Canning et al., 2018; Guo et al., 2017; Lee and Gleeson, 2011; Louvi and Grove, 2011; Marley and von Zastrow, 2012). Despite the emerging role of primary cilia in neurodevelopmental disorders, their participation in the underlying disease mechanisms remains largely unexplored. In particular, a better understanding of their involvement in physiological human brain development is required.

Sonic hedgehog (SHH) signalling is a key factor in normal and abnormal nervous system development. Crucially, its pathway activity is controlled by primary cilia. In the absence of SHH protein, ciliary expression levels of SMO, the main cellular transducer of Hedgehog signals, are low since PTCH1 prevents SMO from entering the cilium (Rohatgi et al., 2007). In addition, activation of Protein Kinase A (PKA) through the negative regulator GPR161 (Mukhopadhyay et al., 2013) results in the phosphorylation of the GLI3 transcription factor and its proteolytic processing at the base of the cilium. As a result, GLI3 repressor (GLI3R) is formed (Haycraft et al., 2005) that inhibits the expression of Hedgehog target genes. In contrast, upon pathway activation, SHH binds to and represses PTCH1 so that SMO together with SUFU, GLI2 and GLI3 accumulate at the ciliary tip (Corbit et al., 2005; Haycraft et al., 2005; Kopinke et al., 2020; Liem et al., 2009; Rohatgi et al., 2007; Wen et al., 2010). There, the GLI proteins are converted through phosphorylation events into transcriptional activators (GLIA) (Han et al., 2019) that translocate to the nucleus and stimulate the transcription of SHH target genes. Thus, primary cilia are critical for both producing GLI3R and for activating the full length GLI proteins. These activities make cilia ideal candidates for controlling human nervous system development, a function that has hitherto not been examined in detail in human model systems.

Due to their involvement in cell-cell signalling in general and in SHH signalling in particular, primary cilia are likely to be crucial in controlling dorsal/ventral (D/V) patterning of the telencephalon (Andreu-Cervera et al., 2021; Liu et al., 2021; Park et al., 2019). Indeed, several ciliary mouse mutants show a variety of telencephalic patterning phenotypes ranging from extensive ventralisation of the dorsal telencephalon to milder defects in properly forming the pallial/subpallial boundary that separates the dorsal from the ventral telencephalon (Ashique et al., 2009; Besse et al., 2011; Hasenpusch-Theil et al., 2020; Stottmann et al., 2009; Willaredt et al., 2008). In contrast, it has not been investigated whether cilia play a similar role in human cortical development. To address this question, we became interested in the ciliary gene *INPP5E* that is mutated in MORM (mental retardation, obesity, retinal dystrophy and micropenis) syndrome (Jacoby et al., 2009) and in Joubert Syndrome (JS) (Bielas et al., 2009), a ciliopathy characterised by cerebellar defects and malformations of the cerebral cortex in a subset of patients. *INPP5E* encodes Inositol Polyphosphate-5-Phosphatase E, a ciliary membrane associated protein that hydrolyses the phosphatidylinositol polyphosphates PI(4,5)P_2_ and PI(3,4,5)P_3_ (Bielas et al., 2009; Jacoby et al., 2009). In this way, INPP5E regulates the phosphoinositol composition of the ciliary membrane and creates a specific phosphoinositide distribution that has been linked to protein trafficking and Hedgehog signalling to either repress or activate the pathway in a tissue specific manner (Chavez et al., 2015; Constable et al., 2020; Garcia-Gonzalo et al., 2015; Hasenpusch-Theil et al., 2020). *Inpp5e* also regulates cilia assembly (Xu et al., 2016), stability (Jacoby et al., 2009) and disassembly (Plotnikova et al., 2015) by controlling primary cilia excision (Phua et al., 2017). In addition to these cellular functions, *Inpp5e* knock-out and conditional mouse mutants recapitulate many of the defects observed in human JS patients (Hakim et al., 2016; Hasenpusch-Theil et al., 2020; Jacoby et al., 2009; Ukhanov et al., 2021). In particular, we recently showed that loss of *Inpp5e* function leads to a defective pallial/subpallial boundary and has profound effects on cortical stem cell functions (Hasenpusch-Theil et al., 2020).

Based on these widespread functions in ciliary biology and in mammalian neural and non-neural development, *INPP5E* presents an excellent candidate to study potential functions of primary cilia in human corticogenesis. Using CRISPR/CAS9 mutagenesis, we engineered human induced pluripotent stem cells (iPSCs) to carry an *INPP5E* loss-of-function mutation and generated cortical organoids from these lines. We show that inactivating *INPP5E* resulted in ventralised organoids that formed ventral telencephalic progenitors and neurons rather than cortical projection neurons. The mutation also caused an up-regulation of SHH signalling which was necessary and sufficient for this ventralisation. Mechanistically, we show that the *INPP5E* mutation interfered with the function of the ciliary transition zone and led to an increased accumulation of SMO in the cilium.

## RESULTS

### Primary cilia in the developing human telencephalon

Cell-cell signalling mediated by primary cilia plays crucial roles in murine corticogenesis (Andreu-Cervera et al., 2021; Hasenpusch-Theil and Theil, 2021; Liu et al., 2021). As a first step to establish ciliary function in human cortical development, we investigated the presence of primary cilia in the developing human telencephalon. We immunostained the forebrain of a PCW 8 human embryo to reveal its D/V subdivisions. At this stage, SOX2 was expressed in progenitor cells in the ventricular zone (VZ) throughout the telencephalon with some scattered SOX2+ progenitors in the forming cortical subventricular zone (SVZ) (Fig. 1A, D). PAX6 expression was confined to the dorsal telencephalon where the protein was detected in apical radial glial cells (aRGCs) in the VZ and in a few progenitors in the SVZ with a lateral high to medial low expression gradient (Fig, 1B, E) as described previously (Clowry et al., 2018; Qin et al., 2020). Moreover, cortical projection neurons residing in the forming cortical plate were identified by the expression of TBR1 (Fig. 1C, F). In contrast, VZ progenitor cells of the lateral ganglionic eminence expressed GSX2 and DLX2 that was also detected in the SVZ (Fig. 1G, H, J, K). Finally, OLIG2 is expressed in the MGE and LGE progenitor domains and in scattered OLIG2+ oligodendrocyte precursor cells (Fig. 1I, L). We next performed immunostainings for some of these regional markers in combination with the axonemal marker ARL13B to reveal the presence of primary cilia. This analysis identified cilia projecting from the apical surface of radial glial cells into the lumen of the lateral ventricle in both the cortex and the LGE (Fig. 1M, O). At this stage, however, cilia were not found on cortical projection neurons (Fig. 1N). Thus, cilia are present on progenitor cells to receive signals important for D/V patterning of the human telencephalon.

**Figure 1:**
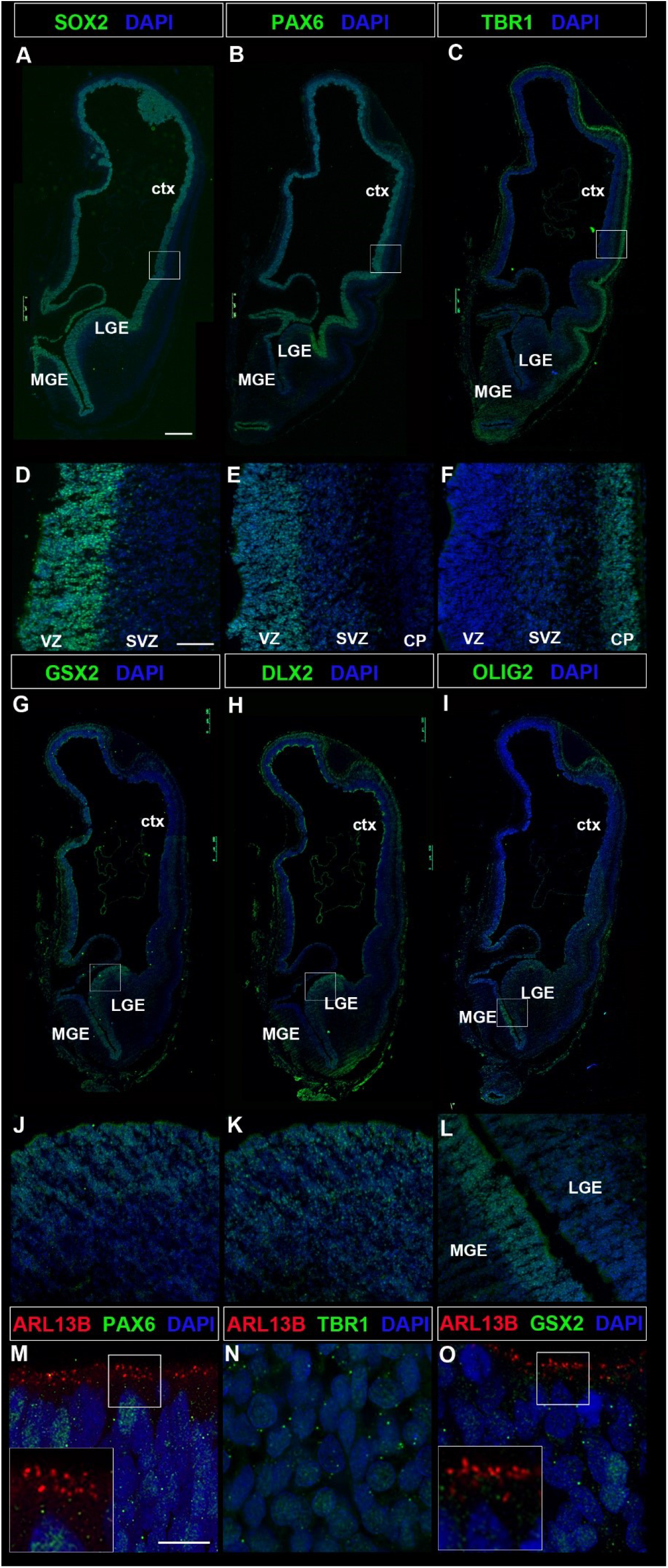
Cilia in the human telencephalon. Coronal sections of the telencephalon of a human 8 weeks old embryo immunostained with the indicated markers. (A, D) SOX2 expression in dorsal and ventral telencephalic progenitors. (B, C, E, F) PAX6 and TBR1 expression are confined to dorsal telencephalic progenitors and neurons, respectively. (G-L) GSX2 (G, J), DLX2 (H, K) and OLIG2 (I, L) expression are confined to the ventral telencephalon. (M-O) ARL13B+ cilia projecting from progenitor cells into the ventricular lumen in the dorsal (M) and ventral (O) telencephalon. Note the absence of ARL13B expression in TBR1+ projection neurons (N). ctx, cortex; MGE, medial ganglionic eminence; LGE, lateral ganglionic eminence; SVZ, subventricular zone; VZ, ventricular zone. Scale bars: 500 μm (A), 100 μm (D), 2.5 μm (M).

### Generation and initial characterisation of *INPP5E* mutant iPSCs

To test the role of primary cilia in telencephalon development, we started to investigate the function of the *INPP5E* gene which is essential for cilia mediated signalling (Bielas et al., 2009; Chavez et al., 2015; Garcia-Gonzalo et al., 2015; Jacoby et al., 2009) and which is required in embryonic and adult neural stem cells (Chavez et al., 2015; Hasenpusch-Theil et al., 2020). To this end, we generated mutant human iPSC cell lines with a homozygous D477N mutation (Fig. 2A) using a clustered regularly interspaced short palindromic repeats (CRISPR)/Cas9 approach. This mutation represents an enzymatic null mutation and has been widely used to characterize *INPP5E* function (Kong et al., 2006). A guide RNA (gRNA) was selected that had at least three mismatches to potential off-target sites which are highly homologous to the on-target site. The gRNA/Cas9 plasmid was co-transfected with a template oligonucleotide carrying the D477N mutation into control iPSCs (Johnstone et al., 2019; Selvaraj et al., 2018; Vasistha et al., 2019). In this way, we achieved a ca. 8% mutation efficiency and identified 23 clones with the desired homozygous mutation as confirmed by restriction fragment length polymorphism and Sanger sequencing (Fig. 2B). Three of these clones (1C2, 2A6 and 2G2) were chosen for further analyses. These mutant clones were karyotypically normal and retained the pluripotency markers NANOG, OCT3/4 and TRA-1-60 (Supplementary Figure 1). In all three clones, we did not detect off-target activity of the gRNA for the six highest candidate off-target sites as assessed by Sanger sequencing.

**Figure 2:**
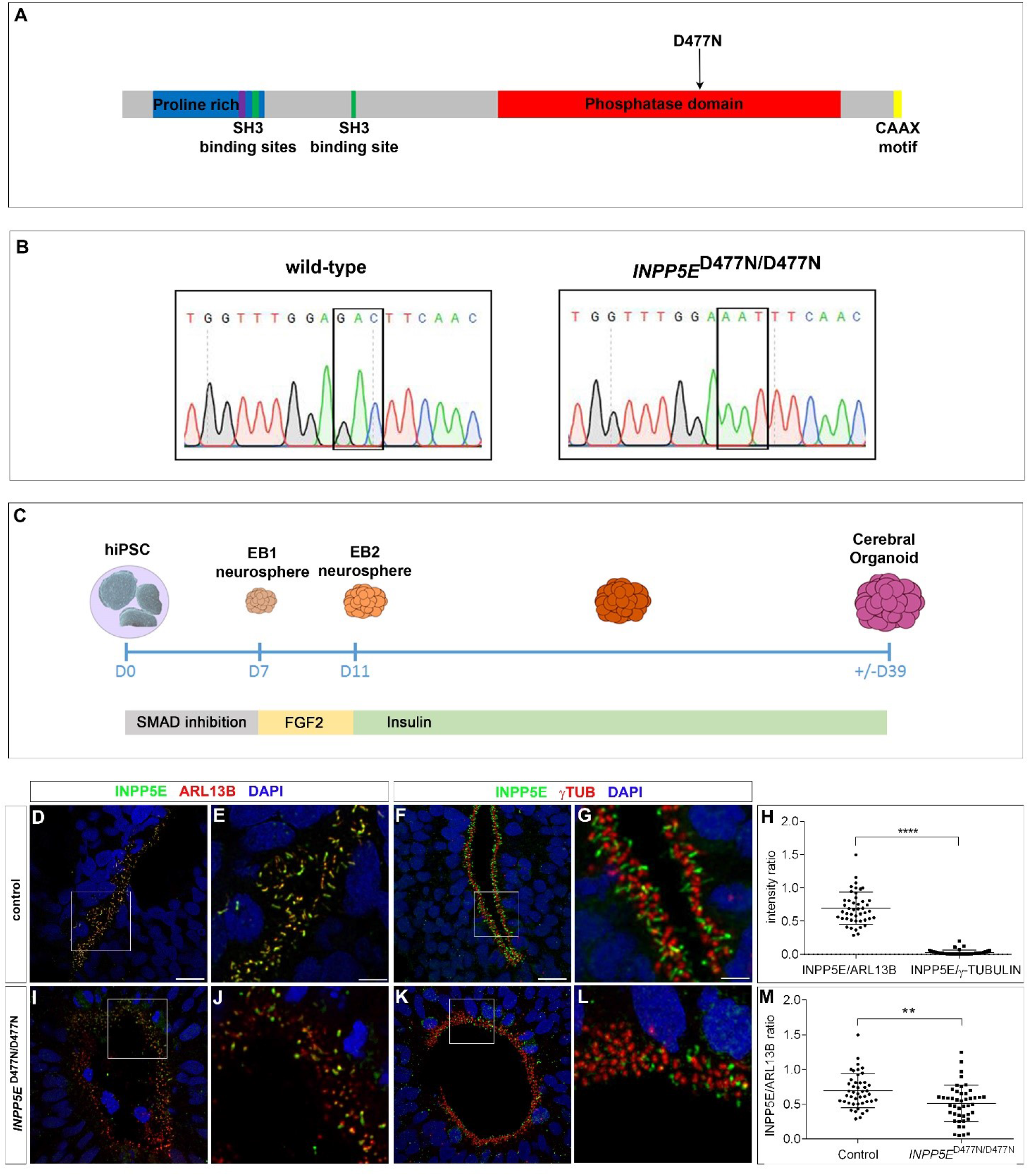
*INPP5E* mutagenesis and expression in cortical organoids. (A) Schematic domain structure of the INPP5E protein, the D477N mutation affecting INPP5E’s phosphatase activity is indicated. (B) Sequencing traces confirming successful mutagenesis. (C) Schematic of the protocol used to generate cortical organoids. (D-M) INPP5E expression in cilia of control (D-G) and in *INPP5E*^D477N/D477N^ organoids (I-L). INPP5E protein was confined to the axoneme and excluded from the basal body while expression levels were reduced in *INPP5E*^D477N/D477N^ organoids. (H, M) Quantification of immunostainings. Statistical data are presented as means ± 95% confidence intervals (CI); Mann-Whitney tests; n = 45 cilia from three different lines; ** p < 0.01; **** p<0.0001. Scale bar: 10 μm (E, G), 2.5 μm (D, F).

3 control and 3 *INPP5E* mutant iPSC lines were differentiated into cerebral organoids using a modified Lancaster protocol (Fig. 2C) (Lancaster et al., 2013). After dual-Smad inhibition stimulating neural induction and embryoid body (EB) formation, FGF2 was added to the culture medium to promote neuroepithelial expansion. EBs were maintained on a shaking incubator to enhance oxygen exchange and nutrient absorption. At week 4, control and mutant organoids developed large neuroepithelial loops that increased over the next couple of weeks before cerebral organoids were harvested (Day 39). We first characterised the presence of primary cilia in these organoids. Control organoids form large, elongated neuroepithelia whereas mutant organoids often formed smaller, rosette-like structures. ARL13B and γ TUB staining labelling the ciliary axoneme and the basal body, respectively, revealed primary cilia emanating from apical radial glia cells into the lumen in organoids of both genotypes as found in human fetal cortical tissue (Fig. 2D, F, I, K). We next examined INPP5E protein expression. In control organoids, INPP5E protein was confined to the ciliary axoneme and excluded from the basal body. This pattern was maintained in mutant organoids, though the expression of the INPP5E^D477N/D477N^ mutant protein was reduced (Fig. 2D-M). Taken together, these findings indicate that control and *INPP5E*^D477N/D477N^ mutant cortical organoids establish a correct organisation of the neuroepithelium with respect to the apical location of the primary cilium projecting into the lumen and a restricted expression of the INPP5E protein to the cilium.

### *INPP5E*^D477N/D477N^ organoids are ventralised

After having established neural differentiation, we investigated the type of neuroepithelium formed in control and *INPP5E*^D477N/D477N^ mutant organoids. *FOXG1* encodes a transcription factor expressed throughout the telencephalon (Xuan et al., 1995). We detected strong FOXG1 expression in neural progenitors and neurons of both types of organoids consistent with the idea that the protocol generated organoids with telencephalic identity (Fig. 3A, B). After its establishment, the telencephalon becomes subdivided into dorsal and ventral regions that give rise to the cerebral cortex and the ganglionic eminences, respectively. To determine the regional telencephalic identity of the organoids, we performed immunofluorescence analyses with various dorsal and ventral specific markers. PAX6 and EMX1 label aRGCs in the developing cortex. In contrast to control organoids, these markers were hardly expressed in *INPP5E*^D477N/D477N^ organoids (Fig. 3C-F). During cortical neurogenesis, aRGCs undergo asymmetric cell divisions to form cortical neurons directly or indirectly via the production of basal progenitors. Basal progenitors and early born cortical neurons are characterized by the expression of TBR2 and TBR1/CTIP2, respectively (Englund et al., 2005). TBR2+, TBR1+ and CTIP2+ cells were readily identified in control organoids where they separate into different layers, the subventricular zone and cortical plate, respectively, on top of the ventricular zone in a similar arrangement to that found during corticogenesis (Fig. 3G, I, K). Interestingly, TBR2, TBR1 and CTIP2 expression were largely absent from mutant organoids (Fig. 3H, J, L) suggesting that the *INPP5E*^D477N/D477N^ mutation interferes with the formation of major dorsal telencephalic cell types.

**Figure 3:**
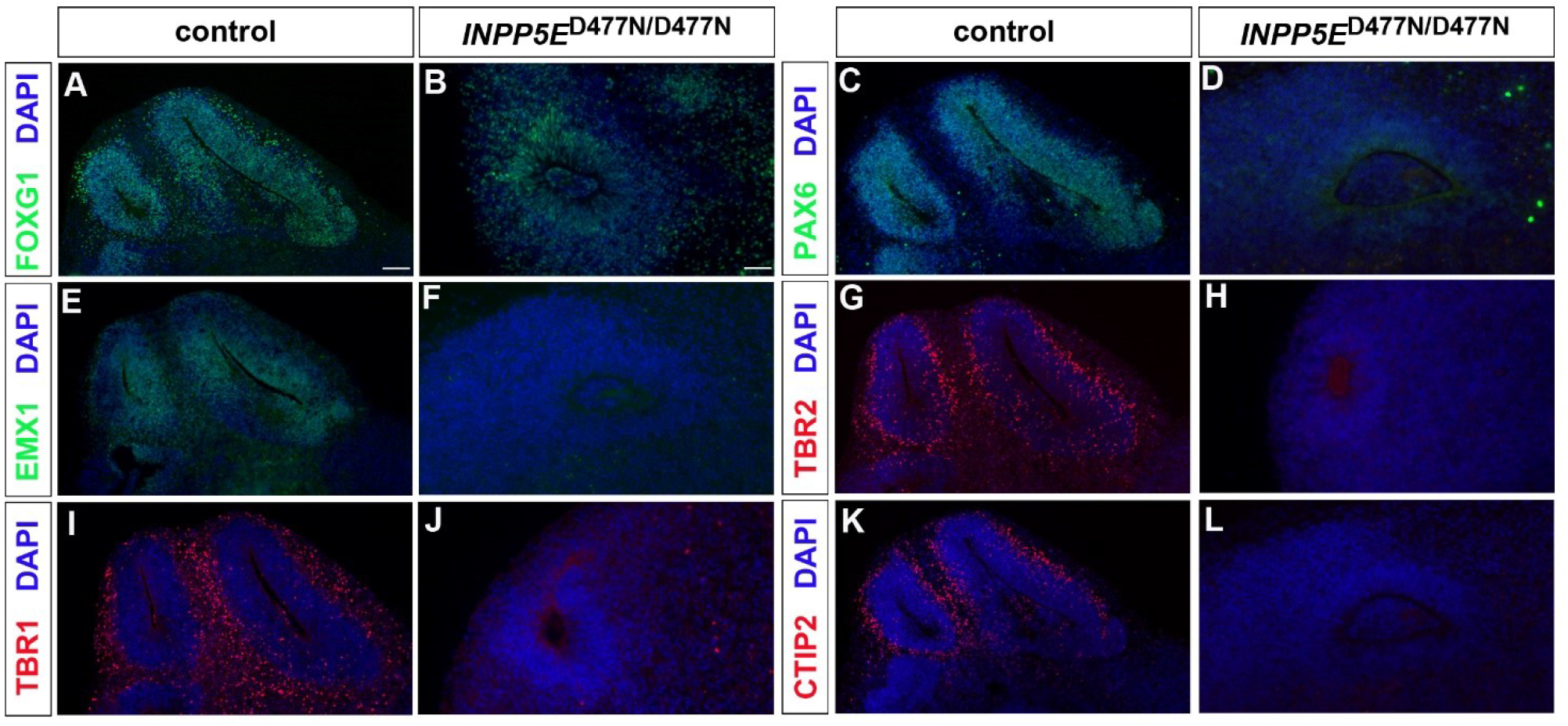
The *INPP5E* mutation interferes with dorsal telencephalic marker gene expression. Control and *INPP5E*^D477N/D477N^ organoids were immunostained with the indicated markers. (A, B) The telencephalon marker FOXG1 was expressed in organoids of both markers. (C-F) The dorsal aRGC markers PAX6 (C, D) and EMX1 (E, F) were expressed in control but not in mutant organoids. (G, H) TBR2 labelled basal progenitor cells which are largely absent in *INPP5E*^D477N/D477N^ organoids. (I-L) Expression of TBR1 and CTIP2, characteristic of deep layer cortical projection neurons, is largely absent in mutant organoids. Scale bars: 100 μm (A), 50 μm (D)

The expression of FOXG1 taken together with the lack of dorsal marker protein expression raise the possibility that *INPP5E* mutant organoids form ventral telencephalic structures. To test this idea, we systematically investigated the formation of ventral progenitor cells and their neuronal derivatives in *INPP5E* mutant organoids. In embryonic development, the ventral telencephalon contains GSX2+ and DLX2+ progenitor cells. In addition, it generates OLIG2+ oligodendrocyte precursor cells (OPCs). In mouse embryogenesis, these are formed earlier in the ventral telencephalon compared to the cortex where OPC formation is delayed (Kessaris et al., 2006). A similar temporal order of OPC formation was recently observed in human dorsal and ventral organoids (Kim et al., 2019). In *INPP5E*^D477N/D477N^ mutant organoids, GSX2+, DLX2+ and OLIG2+ progenitors were found abundantly whereas controls only contained a few isolated cells (Fig. 4A-F). Subsequently, we determined the expression of markers characteristic of ventral telencephalic neurons. Striatal cholinergic interneurons arising from the lateral ganglionic eminence express the ISL1 LIM homeodomain transcription factor which is required for their survival and differentiation (Elshatory and Gan, 2008; Wang and Liu, 2001). Immunofluorescence analysis revealed the widespread presence of ISL1/2+ neurons surrounding neuroepithelial structures in mutant but not in control organoids (Fig. 4E, F). The MGE and CGE also give rise to GABAergic interneurons that migrate from their birth place in the ventral telencephalon to the cortex where they eventually settle in the cortical plate. MGE derived interneurons are characterized by NKX2.1 or Somatostatin (SST) expression while CGE interneurons are positive for COUP-TFII. Neurons expressing these markers were found in *INPP5E*^D477N/D477N^ organoids (Fig. 4G-L). Thus, mutant organoids lack dorsal telencephalic cell types but express a number of ventral telencephalic progenitor and neuron markers. This expression profile is consistent with the idea that the *INPP5E*^D477N/D477N^ mutation leads to a ventralisation of cortical organoids.

**Figure 4:**
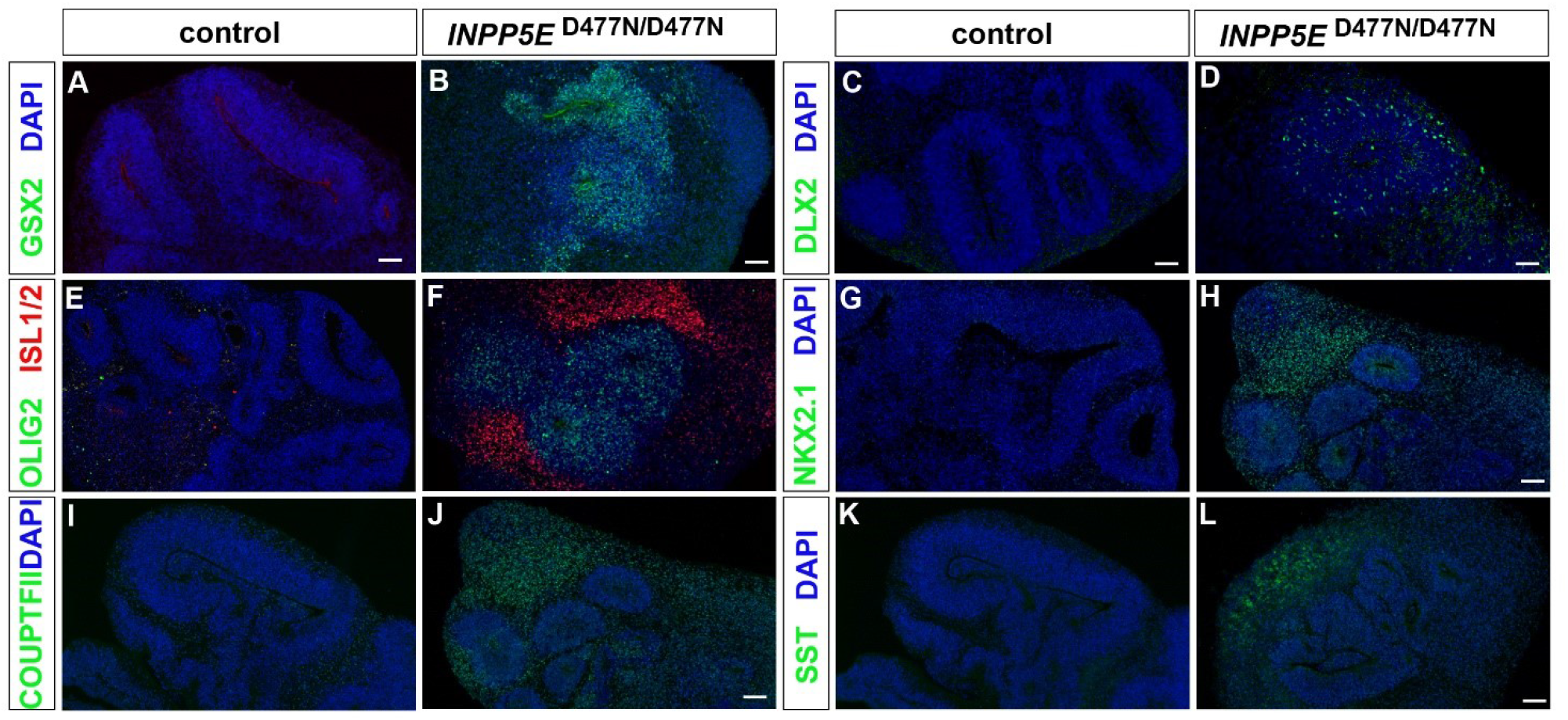
*INPP5E*^D477N/D477N^ organoids are ventralised. Immunofluorescence analyses of control and *INPP5E*^D477N/D477N^ organoids with the indicated markers. (A-F) Expression of the ventral telencephalic progenitor markers GSX2 (A, B), DLX2 (C, D) and OLIG2 (E, F) in mutant but not in control organoids. Similarly, expression of ISL1/2 characteristic of spiny medium interneurons is detected in *INPP5E* mutant organoids only (E, F). (G-L) Expression analyses of the ventral telencephalic interneuron markers NKX2.1 (G, H), COUPTFII (I, J) and SST (K, L). Scale bars: 100 μm (A, CH, J, L), 50 μm (B, D).

### Sonic hedgehog signalling is up-regulated in *INPP5E*^D477N/D477N^ organoids

Next, we aimed at identifying the mechanisms behind this ventralisation. It has been well established that several signalling molecules are crucial for D/V patterning of the telencephalon (Rallu et al., 2002a). Most notably, SHH promotes ventral telencephalic development and represses dorsal cell fate acquisition in mice (Rallu et al., 2002b) and human organoids (Bagley et al., 2017; Birey et al., 2017). Given that cilia are important regulators of SHH signalling we hypothesized that the ventralised phenotype of *INPP5E*^D477N/D477N^ organoids originated from an up-regulation of SHH signalling. To test this hypothesis, we first examined the expression of the SHH target genes *PTCH1* and *GLI1*. Using in situ hybridisation we found expression of both markers in the neuroepithelial rosettes of the mutant organoids but not in the ventricular zones of control organoids (Fig. 5A, B, D, E). We confirmed this finding in quantitative RT-PCR experiments that showed a 4.8fold and 3.1fold increase in *PTCH1* and *GLI1* expression, respectively (Fig. 5C, F). In addition to their role in activating the GLI transcription factors in the presence of SHH, cilia play an important role in processing the GLI3 full length protein (GLI3FL) to form the GLI3 repressor (GLI3R) which predominates in the dorsal telencephalon of mice (Fotaki et al., 2006). Western blot analyses of organoid protein extracts revealed decreased GLI3R expression levels (Fig. 5G, H) while GLI3FL levels and the GLI3FL/GLI3R ratio were not affected (Fig. 5I, J). Taken together, these data show an increase in SHH signalling coinciding with a reduced formation of GLI3R in *INPP5E*^D477N/D477N^organoids.

**Figure 5:**
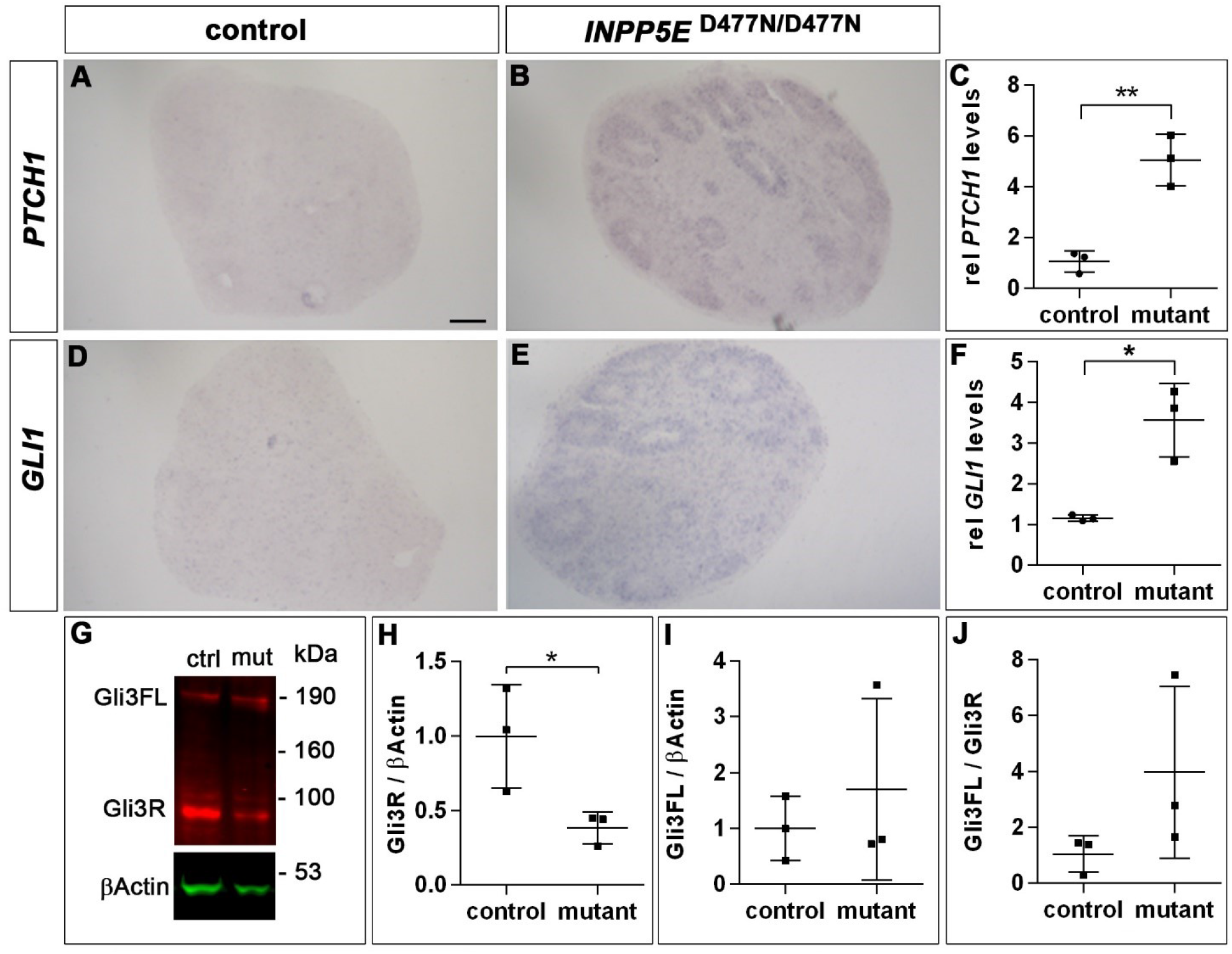
SHH signalling is up-regulated in *INPP5E*^D477N/D477N^ organoids. (A, B, D, E) in situ hybridisation to detect *PTCH1* (A, B) and *GLI1* (D, E) expression in control (A, D) and *INPP5E* mutant organoids (B, E). (C, F) Representative example of qRT-PCR analyses showing increased *PTCH1* (C) and *GLI1* (F) mRNA expression levels relative to *ATP5*. (G) GLI3 Western blot on organoid tissue revealed the GLI3 full length (FL) and repressor (R) forms. (H-J) Quantification of GLI3 Western blot. GLI3R levels are decreased in *INPP5E*^D477N/D477N^ organoids (H), while GLI3FL levels (I) and the GLI3R/GLI3FL ratio (J) are not affected. All statistical data are presented as means ± SD; unpaired t-tests (n=3) (C, H-J) and unpaired t-tests with Welch’s correction (F); * p < 0.05; ** p < 0.01. Scale bar: 100 μm (A).

### SHH signalling is necessary and sufficient to ventralise cortical organoids

We next tested the role of the up-regulation of SHH signalling in the induction of ventral marker gene expression in *INPP5E*^D477N/D477N^ mutant organoids. To this end, we repeated the organoid experiment but added the SHH antagonist cyclopamine to the culture medium on Day 7 for the remainder of the organoid culture to block the constitutive SHH signalling caused by the *INPP5E* mutation (Fig. 6A). In a separate experiment, we also investigated whether activating SHH signalling was sufficient to induce a ventralisation in cortical organoids derived from control iPSCs. To this end, we treated organoids after neural induction from Day 7 for one week with the SHH agonist Purmorphamine (Fig. 6A). In previous experiments, we used this treatment for the generation of ventral telencephalon derived oligodendrocytes from human iPSCs (James et al., 2021; Livesey et al., 2016).

**Figure 6:**
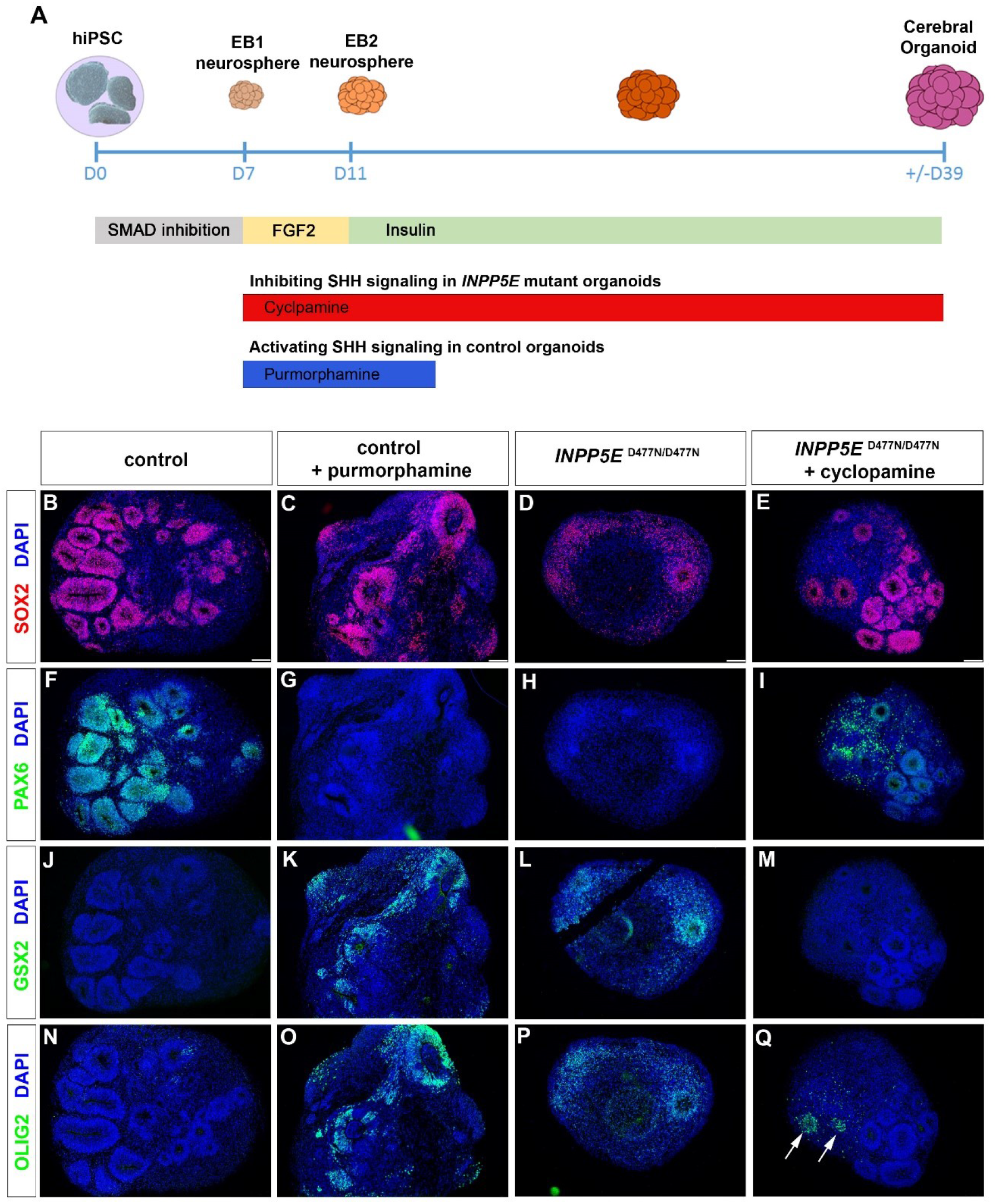
SHH signalling is necessary and sufficient to ventralise cortical organoids. (A) Experimental protocol to either block or ectopically activate SHH signalling in cortical organoid cultures by the addition of Cyclopamine and Purmorphamine, respectively. (B-Q) Organoids stained with the indicated antibodies after treatment with Purmorphamine (C, G, K, O) and Cyclopamine (E, I, M, Q). (B-E) SOX2 expression revealed neuroepithelia. (F-I) PAX6 expression is reduced in control organoids treated with Purmorphamine but up-regulated after inhibiting SHH signalling as compared to untreated *INPP5E* mutant organoids. (J-Q) GSX2 and OLIG2 expression occurs in only a few cells in control organoids (J, N) but is strongly up-regulated after activation of SHH signalling (K, O) or in untreated *INPP5E* mutant organoids (L, P). (M, Q) Treating mutant organoids with Cyclopamine inhibits GSX2 and OLIG2 expression except for a few cells (arrows in Q). Scale bars: 100 μm (A-D).

Day 39 organoids from both experiments were harvested and stained for SOX2 to identify neuroepithelial structures (Fig. 6B-E) and for various dorsal and ventral telencephalic progenitor markers. These analyses first confirmed the expression of PAX6 and the absence of the ventral markers GSX2 and OLIG2 in control untreated organoids (Fig. 6F,J, N). In contrast, these ventral markers were up-regulated with a concomitant down-regulation of PAX6 in control organoids treated with Purmorphamine (Fig. 6G, K, O) suggesting that activation of SHH signalling was sufficient to ventralise cortical organoids. As described above *INPP5E*^D477N/D477N^ mutant organoids raised under control conditions also lacked PAX6 expression and showed an up-regulation of GSX2, and OLIG2 (Fig. 6H, L, P). Cyclopamine treatment, however, led to a mild up-regulation of neuroepithelial PAX6 (Fig. 6I). Moreover, GSX2 and OLIG2 expression were largely absent with a few neuroepithelial rosettes still expressing OLIG2 (Fig. 6M, Q). Taken together, this staining pattern indicates that cyclopamine treatment confers a partial rescue of the ventralisation caused by the *INPP5E* mutation.

### TULP3 and GPR161 are enriched in the *INPP5E*^D477N/D477N^ mutant cilium

Next, we aimed to identify the molecular mechanisms that led to perturbed SHH signalling in the *INPP5E* mutant organoids. The INPP5E enzyme hydrolyses the 5-phosphate from PI(4,5)P2 to produce PI(4)P. Consistent with INPP5E’s axonemal localisation, PI(4)P levels were recently shown to be enhanced in the axoneme while PI(4,5)P_2_ was limited to the ciliary base (Chavez et al., 2015; Garcia-Gonzalo et al., 2015). Conversely, loss of *INPP5E* function resulted in PI(4,5)P_2_ accumulation in the axonemal membrane and at the ciliary tip and to altered Hedgehog signalling (Chavez et al., 2015; Constable et al., 2020; Garcia-Gonzalo et al., 2015). Direct detection and distinction of phosphoinositides is technically difficult as their fluidity and lipophilic characteristics pose problems for fixation and staining. Therefore, we stained for the PI(4,5)P_2_ binding protein TULP3, previously shown to be increased in *Inpp5e*-deficient cilia (Chavez et al., 2015; Constable et al., 2020; Garcia-Gonzalo et al., 2015), as an indirect read-out of PI(4,5)P_2_ distribution. As the previously described change in phosphoinositide localisation caused by *INPP5E* mutations primarily affected the axoneme, we studied the proportion of TULP3+ cilia and TULP3 expression levels in the axoneme by double immunofluorescence for TULP3 and the axonemal marker ARL13B. In control organoids, a small proportion of cilia were positive for TULP3 but TULP3 expression was detected in the majority of *INPP5E*^D477N/D477N^ mutant cilia (Fig. 7A-C). Moreover, comparison of TULP3/ARL13B intensity ratios showed a significant increase in *INPP5E*^D477N/D477N^ organoids compared to controls (Figure 7D).

**Figure 7:**
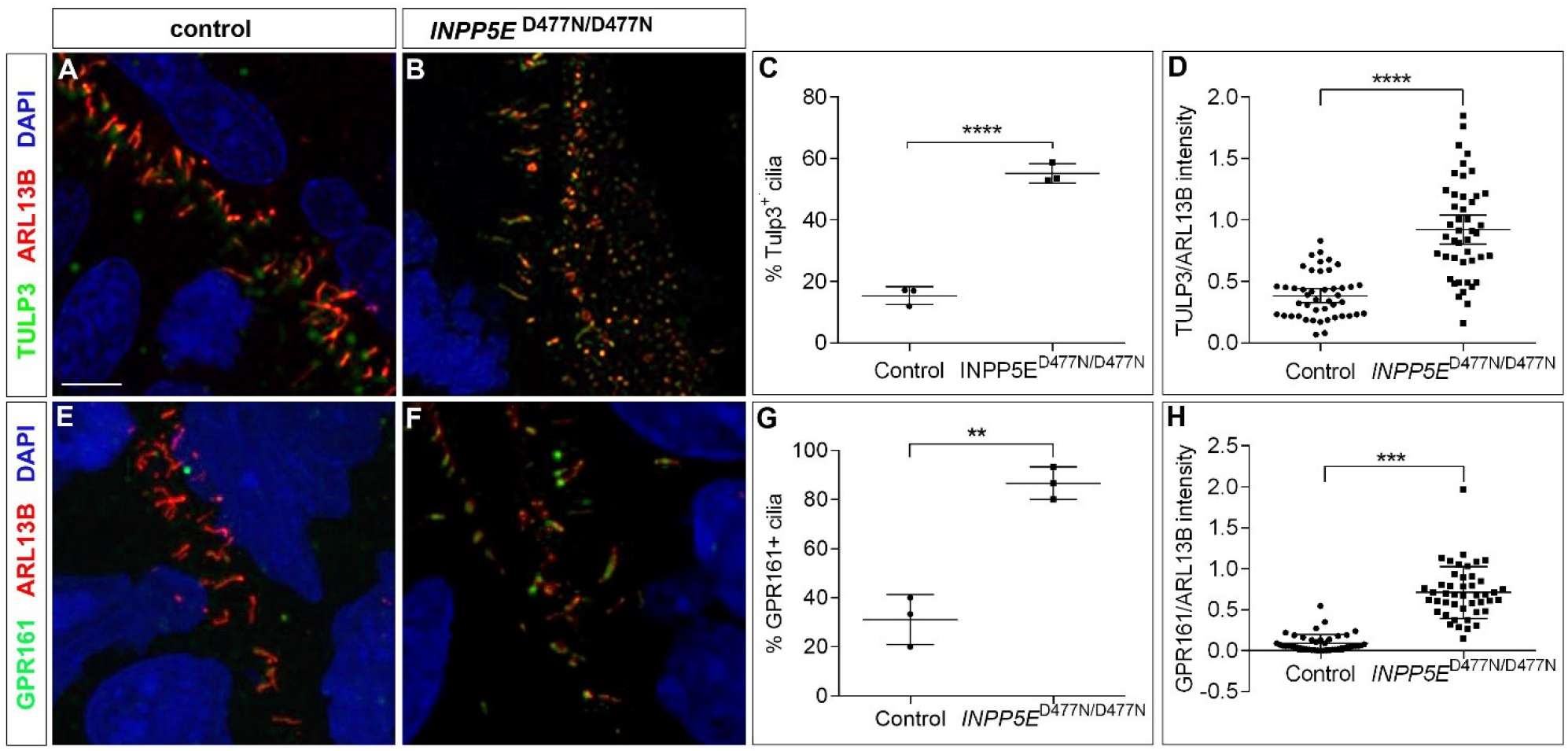
Ciliary localization of TULP3 and GPR161. (A, B, E, F) Immunofluorescence analyses of TULP3 (A, B) and GPR161 (C, D) ciliary expression in control (A, E) and *INPP5E*^D477N/D477N^ organoids (B, F). (C, D, G, H) Quantification of immunostainings. (A-D) TULP3 expression in the axoneme was up-regulated and found in most *INPP5E* mutant cilia but not in control cilia. (E-H) GPR161 was expressed at higher levels in almost all cilia in *INPP5E*^D477N/D477N^ organoids. Statistical data are presented as means ± 95% confidence intervals (CI); unpaired t-tests (C, G), unpaired t-test with Welch’s correction (D) and Mann Whitney test (H); n=3 lines for (C, G) and n = 45 cilia from three different lines (D, H); ** p<0.01; *** p < 0.001; **** p<0.0001. Scale bar: 2.5 μm.

As TULP3 recruits the SHH antagonist GPR161 to the cilium (Chavez et al., 2015; Garcia-Gonzalo et al., 2015), we determined whether the increase in TULP3 in *INPP5E*^D477N/D477N^ organoids coincided with an enrichment of GPR161 in the cilium. Double immunofluorescence labelling for ARL13B and GPR161 revealed an increase in the proportion of GPR161+ cilia and a higher GPR161/ARL13B intensity ratio in *INPP5E* mutant organoids as compared to control organoids (Figure 7E-H). Taken together, these findings indicate an enrichment of TULP3 and GPR161 in the *INPP5E*^D477N/D477N^ axoneme. This is consistent with an higher amount of PI(4,5)P_2_ in the axonemal membrane due to an impairment of INPP5E phosphatase activity.

### Increased ciliary expression of SMO in *INPP5E*^D477N/D477N^ organoids

To examine the effect of INPP5E on SHH signalling, we investigated the expression and localisation of SHH signalling components. SMO is the main cellular transducer of hedgehog signals. In the absence of hedgehog proteins, SMO’s ciliary levels are low since PTCH1 prevents it from entering the cilium (Rohatgi et al., 2007). Upon pathway activation, SHH binds to and inhibits PTCH so that SMO accumulates in the cilium where it initiates the downstream signalling cascade (Corbit et al., 2005; Rohatgi et al., 2007). To compare SMO expression in the cilium, organoids were double stained for SMO and ARL13B. This revealed a significantly increased proportion of SMO+ cilia and SMO/ARL13B intensity ratios in *INPP5E*^D477N/D477N^ organoids (Fig. 8A-D). These enriched SMO levels in the cilium are a strong indicator for enhanced SHH signalling in the *INPP5E*^D477N/D477N^ lines. Following this result, we studied ciliary expression of SUFU and of GLI2, the main repressor and transcriptional activator of SHH signalling, respectively. Upon pathway activation, both proteins enter the cilium and accumulate at the ciliary tip (Corbit et al., 2005; Haycraft et al., 2005; Wen et al., 2010). Given these well characterized changes in the distribution of SUFU and GLI2 proteins we determined their localisation in control and *INPP5E*^D477N/D477N^ organoids. For both markers, however, we did not find significant changes either in the proportion of positive cilia or in their expression levels (Fig. 8E-L). Collectively, these data indicate an enrichment of SMO in the cilium of *INPP5* mutant RGCs but no change for SUFU and GLI2.

**Figure 8:**
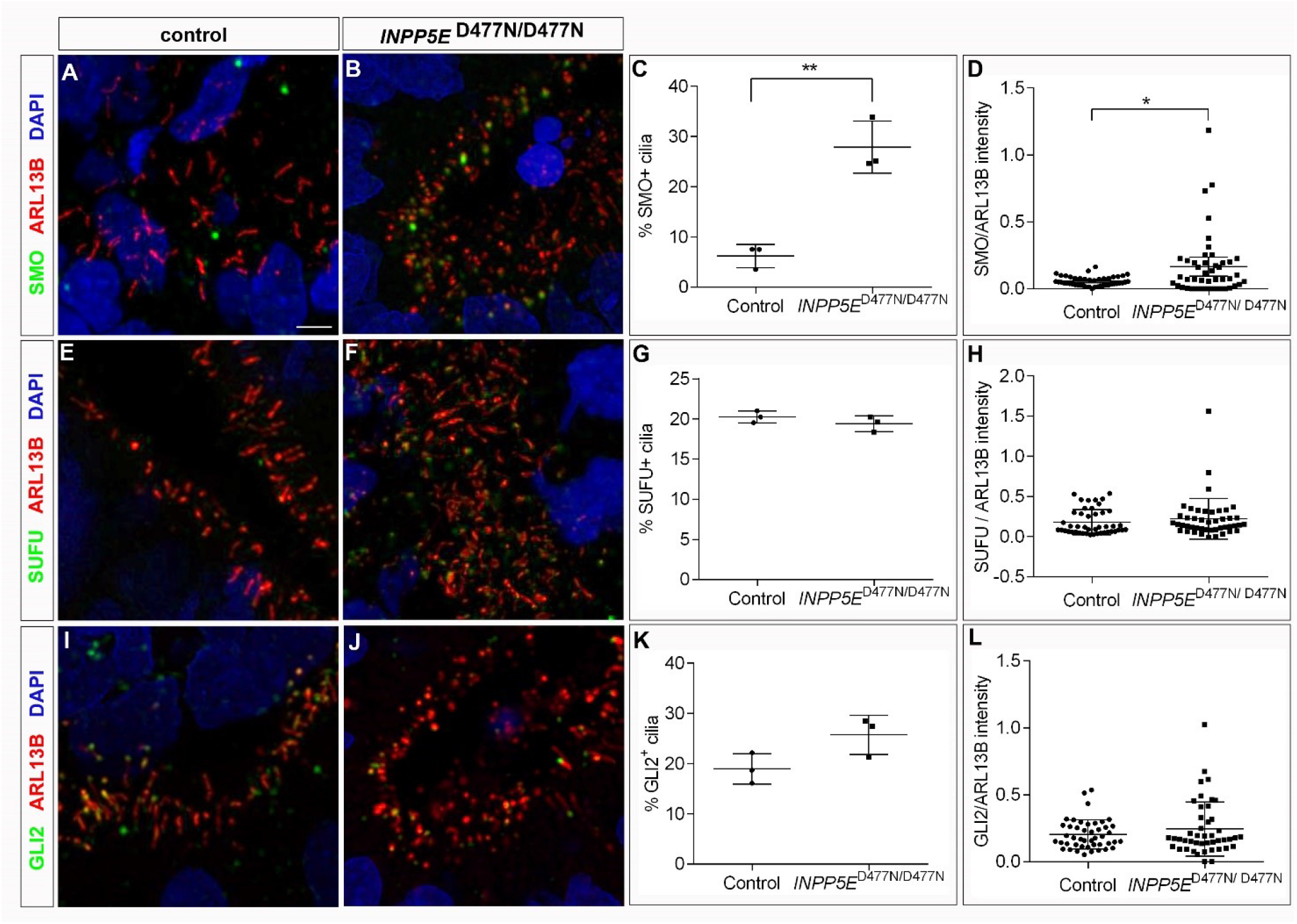
Ciliary expression of SHH signalling components. Immunofluorescence analyses of SMO (A, B), SUFU (E, F) and GLI2 (I, J) ciliary expression in control (A, E, I) and *INPP5E*^D477N/D477N^ organoids (B, F, J). (C, D, G, H, K, L) Quantification of immunostainings. (A-D) SMO was expressed in a higher proportion of cilia and at higher levels in *INPP5E*^D477N/D477N^ organoids. (E-L) There were no significant changes either in the proportion of positive cilia or in the expression levels for SUFU and GLI2. Statistical data are presented as means ± 95% confidence intervals (CI); unpaired t-test (C, G, K) and Mann Whitney tests (D, H, L); n=3 lines for (C, G, K) and n = 45 cilia from three different lines (D, H, L); * p < 0.05; ** p<0.01. Scale bar: 2.5 μm.

### The *INPP5E*^D477N/D477N^ mutation affects transition zone function

Our expression analysis revealed an enrichment for both positive (SMO) and negative (GPR161) regulators of SHH signalling in the ciliary axoneme of *INPP5E* mutant organoids. To identify the molecular basis for this finding, we became interested in the transition zone (TZ) that controls entry and exit of proteins in and out of the cilium, respectively (Garcia-Gonzalo et al., 2011; Reiter et al., 2012). Interestingly, *Inpp5e* is required for the molecular organization and maturation of the TZ in mice and flies (Dyson et al., 2017; Gupta et al., 2018) suggesting that *INPP5E* might also be necessary for the integrity of the transition zone in human cortical organoids. To address this possibility, we performed immunofluorescence stainings for TMEM67, a component of the MKS complex that localises to the TZ (Garcia-Gonzalo et al., 2011). Our immunolabeling revealed TMEM67 protein at the TZ of aRGC cilia in control organoids with low expression levels in the ciliary axoneme. In contrast, a large proportion of *INPP5E*^D477N/D477N^ mutant cilia lacked TMEM67 at the TZ while the protein accumulated in the axoneme suggesting that TZ formation was impaired (Fig. 9A-D). To test whether other proteins not involved in SHH signalling might also mislocalise, we analysed the expression of the intraflagellar transport protein IFT88. In control cilia, IFT88 was found at high levels at the base of the cilium with lower expression levels in the axoneme. In mutant cilia, however, we detected IFT88 co-expression with ARL13B in the axoneme in an increased proportion of cilia (Fig. 9E-H). These findings indicate that TZ function might be compromised *INPP5E* mutant cilia.

**Figure 9:**
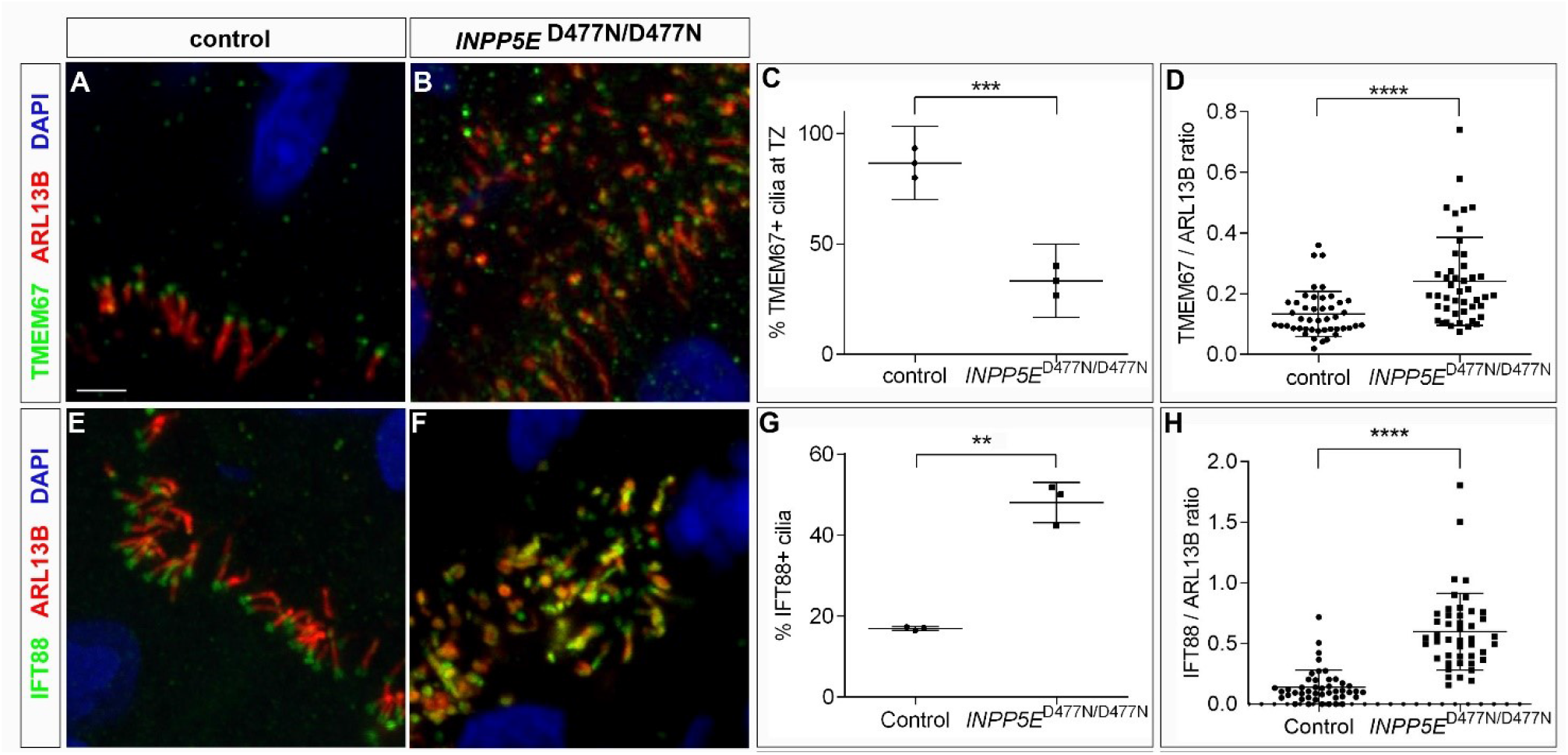
Transition zone in *INPP5E*^D477N/D477N^ organoids. (A, B, E, F) Immunofluorescence analyses of TMEM67 (A, B) and IFT88 (C, D) ciliary expression in control (A, E) and *INPP5E*^D477N/D477N^ organoids (B, F). (A-D) The proportion of cilia with TMEM67 expression at the transition zone was decreased while expression of TMEM67 in the ciliary axoneme was increased in mutant organoids. (E-G) IFT88 was expressed in a higher proportion of ciliary axonemes and at increased levels. Statistical data are presented as means ± 95% confidence intervals (CI); unpaired t-test (C), unpaired t-test with Welch’s correction (G) and Mann Whitney tests (D, H); n = 3 lines (C, G) and n=45 cilia from three different lines for (D, H); ** p < 0.01; *** p < 0.001; **** p<0.0001. Scale bar: 2.5 μm.

## DISCUSSION

Understanding the emerging roles of primary cilia in the pathogenesis of neurodevelopmental disorders requires a better knowledge of their roles in physiological human brain development. Here, we investigated the consequences of inactivating the ciliary *INPP5E* gene on forebrain development in a human cortical organoid model and showed that it is essential for D/V patterning. In the absence of functional *INPP5E*, organoids formed ventral telencephalic progenitors and neurons. This phenotype was caused by an up-regulation of SHH signalling which is necessary and sufficient to induce this ventralisation. The *INPP5E* mutation also led to an accumulation of SMO in the cilium and interfered with the proper formation of the ciliary TZ. Taken together, these findings implicate *INPP5E* as a negative regulator of SHH signalling. They also emphasize the importance of cilia for early human corticogenesis and offer insights into disease mechanism underlying neurodevelopmental disorders.

### Cilia and Dorsal/Ventral Patterning of the human telencephalon

While we are gaining an increasing understanding of mammalian nervous system development from the study of model organisms, this knowledge might not be directly applicable to the developing human brain given its dramatically increased size and its species-specific cellular features (Florio and Huttner, 2014; Hodge et al., 2019). Human organoids represent an excellent experimental system to bridge this knowledge gap (Arlotta and Pasca, 2019; Di Lullo and Kriegstein, 2017). Here, we applied cortical organoid cultures to investigate potential roles of primary cilia in an early step in telencephalic development, namely its subdivision in specific dorsal and ventral territories from which the cortex and basal ganglia develop, respectively. These organoids form large neuroepithelial structures with primary cilia projecting from the apical surface of progenitor cells into the ventricular lumen reflecting the arrangement in the forebrain of a PCW 8 human embryo. Moreover, inactivating the phosphatase activity of *INPP5E* which is critical for ciliary biology and cilia mediated signalling resulted in a striking formation of ventral telencephalic progenitors and neurons due to an activation of SHH signalling. This ventralisation is more severe than the mild defects at the pallial/subpallial boundary we observed in *Inpp5e* mutant mice (Hasenpusch-Theil et al., 2020). The greater severity could be due to an absence of dorsalizing WNT signals produced by the cortical hem (Grove et al., 1998). Alternatively, human and mouse forebrain may follow different paths to establish distinct dorsal and ventral domains in the telencephalon. In an in vitro neural differentiation system, D/V patterning of human forebrain involves subsequent activation/transformation steps whereby neuroectoderm first acquires forebrain dorsal fate by default while the acquisition of ventral telencephalic cell fates requires the repression of dorsal transcription factors such as PAX6 through SHH signalling (Chi et al., 2017). In line with this model, neural organoids exhibit dorsal forebrain identity in the absence of any exogenous signals. Moreover, our findings and that of others that the activation of SHH signalling either genetically through mutating *INPP5E* or pharmacologically through treating organoid cultures with Purmorphamine or SHH agonist (SAG) (Bagley et al., 2017; Birey et al., 2017) induces ventral cell fates in organoids provides further evidence for this hypothesis. In contrast, in the mouse mutual repression between (i) Wnt/Shh signalling and (ii) the Pax6/Nkx2.1 transcription factors underlies the specification of dorsal and ventral cell fates (Gunhaga et al., 2003; Sussel et al., 1999). Regardless of this difference, the ventralisation phenotype in *INPP5E* organoids places primary cilia through their control of SHH signalling at the centre of this patterning process. This finding also has implications for our understanding of neurodevelopmental disorders. A widely regarded hypothesis proposes that an imbalance between excitation and inhibition (E/I imbalance) underlies their phenotypical manifestation (Bourgeron, 2009; Kepecs and Fishell, 2014; Marin, 2012). As a consequence of altered SHH signalling, mutations in ciliary genes could alter the size of the dorsal and ventral progenitor domains and thereby the relative proportions of glutamatergic projection neurons and GABAergic inhibitory neurons produced in these different territories. This scenario provides a novel disease mechanism in addition to previously described roles of primary cilia in controlling the tangential migration of murine cortical interneurons (Baudoin et al., 2012; Higginbotham et al., 2012), their morphology and synaptic connectivity (Guo et al., 2017). Taken together, these studies illustrate several potential ways how defects in primary cilia could affect the E/I balance crucial for proper cortical functioning.

### Mechanism of up-regulated SHH signalling in *INPP5E*^D477N/D477N^ organoids

Besides providing insights into a fundamental process in human forebrain development, our study also shed light on the molecular mechanisms by which *INPP5E* controls SHH signalling. Previous analyses established tissue specific roles for *INPP5E* as a positive or negative regulator of SHH signalling (Chavez et al., 2015; Constable et al., 2020; Garcia-Gonzalo et al., 2011). In murine adult neural stem cells and IMCD3 cells, *Inpp5e* enables Shh signalling by limiting the ciliary levels of Hh inhibitors, including the PI(4,5)P_2_ binding protein Tulp3 and Gpr161 (Chavez et al., 2015; Garcia-Gonzalo et al., 2011). TULP3 and GPR161 levels were also increased after *Inpp5e* inactivation in the developing ventral spinal cord (Constable et al., 2020) and in our organoid system. Interestingly, this increase coincided with augmented SHH signalling, raising mechanistic questions how the *INPP5E* mutations could lead to increased SHH signalling in the presence of the TULP3 and GPR161 repressors. The output of SHH signalling is ultimately determined by the relative levels of GLI activator and repressors. Accordingly, Constable et al., 2020 proposed a decrease in GliR production and a concomitant lower GliR/GliA ratio, but their western blot data did not support this hypothesis, probably due to the use of whole embryo extracts rather than tissue specific samples (Constable et al., 2020). Our human organoid samples, however, consisted largely of cortical tissue and indeed our analysis revealed decreased GLI3R levels in *INPP5E* mutant organoids whereas GLI3FL levels and the GLI3FL/GLI3R ratio were not significantly altered. This suggests that cilia mediated control of GLI3R formation is critical for dorsal telencephalic development in human organoids. This finding is in line with our recent observation on the role of *Inpp5e* in murine corticogenesis and is also consistent with the activation/transformation model proposed for D/V patterning in the human telencephalon (Chi et al., 2017).

This leaves the question how loss of *INPP5E* phosphatase activity leads to defective GLI3 processing. While the accumulation of SMO, the main activator of the pathway, is consistent with increased SHH signalling, cilia also contained augmented levels of the SHH repressor GPR161. Interestingly, GPR161 forms a module with a regulatory subunit of PKA that by amplifying cAMP signals modulates PKA activity but exits the cilium in the presence of SHH (May et al., 2021). Hence, GPR161’s accumulation in the *INPP5E*^D477N/D477N^ cilium could lead to prolonged PKA activation, GLI3 hyperphosphorylation and increased proteolytic degradation. Alternatively, the simultaneous presence of both activators and repressors without SHH pathway stimulation was previously noted (Shinde et al., 2020) and indicates that the mechanisms that control entry and exit of these molecules into and out of the cilium in response to SHH were not operating properly. Abnormal transport could be caused by a defect in the BBSome (Hey et al., 2021). Alternatively, we noted an abnormal localisation of the TZ protein TMEM67 in the axoneme strongly suggesting a defect in forming the TZ and/or in TZ barrier function. A disturbance of TZ function as a consequence of *INPP5E* inactivation would fit into a previously described complex interplay between phosphoinositides, the TZ, and SHH signalling that governs the signalling function of the primary cilium. The INPP5E substrates PIP3 and PI(4,5)P_2_ form discrete pools at the TZ (Conduit et al., 2017; Dyson et al., 2017) and several TZ proteins were found to be aberrantly located in the cilium of *Inpp5e* mutants (Dyson et al., 2017) and contain phosphoinositide binding domains (Garcia-Gonzalo and Reiter, 2012). Similar to *INPP5E* mutations, active SHH signalling causes an increase in PIP3 levels at the TZ thereby reducing cilia stability (Dyson et al., 2017). In addition, TZ proteins have been shown to be required for Gli3 processing (Garcia-Gonzalo et al., 2011; Sang et al., 2011; Thomas et al., 2012; Wang et al., 2017), in particular Rpgrip1l controls the activity of the proteasome at the basal body responsible for proteolytic processing of Gli3 (Gerhardt et al., 2015). Finally, loss of the TZ protein Tctn2 phenocopies the neurogenesis defects in *Inpp5e* mouse mutants (Hasenpusch-Theil et al., 2020). Taken together, these findings indicate that a defect in TZ barrier function represents a likely mechanism leading to abnormal GLI3 processing and ventralisation in *INPP5E*^D477N/D477N^ organoids.

## MATERIAL AND METHODS

### Cell culture

The human pluripotent stem cell-lines used in this study were obtained with full Ethical/Institutional Review Board approval by the University of Edinburgh and validated using standard methods including chromosomal analysis, pluripotency and absence of plasmid integration. The iPSC1 line was described previously (Johnstone et al., 2019; Selvaraj et al., 2018; Vasistha et al., 2019), the additional iPSC control lines hPSC2 (male) and hPSC3 (male) were obtained from Cedars-Sinai. All iPSCs were continuously maintained in Essential 8™ medium (Gibco, ThermoFisher) on Matrigel® (Corning) coated 6-well plates. HEK 293 cells were cultured in DMEM/F-12 (Gibco, ThermoFisher) supplemented with 10% Foetal Calf Serum (FCS) (Gibco, ThermoFisher) and 2 mM L-glutamine (Gibco, ThermoFisher). All cell types were maintained at 37 °C in a 5% CO_2_ atmosphere.

### Gene editing by CRISPR/Cas9 Homology-Directed Repair

Generating the *INPP5E* D447N mutation was performed in the iPSC1 line. To ensure that this line did not contain unknown *INPP5E* mutations, the targeted exon and flanking sequences were sequenced prior to gene editing. gRNAs were designed using an online CRISPR design tool (DESKGEN) and were cloned into the pSpCas9(BB)-2A-Puro (PX459) plasmid (Addgene: #48139). To test gRNA efficiency a T7 endonuclease assay was performed. gRNA constructs were transfected with Lipofectamine 2000 (Invitrogen, ThermoFisher) into HEK 293 cells, which had been seeded 16-24 hours prior to transfections. Cells were harvested 48 hours post-transfection and genomic DNA was extracted using the Wizard® SV Kit (Promega). Genomic targeting efficiency for each gRNA was determined through annealing and digestion with T7 Endonuclease I (NEB: #M0302) of PCR products flanking the INPP5E D447N target site. gRNA 5’-CTGTGCGCCCGCCACTCAGG-3’ was determined as optimal for use in gene-editing. iPSCs at 70-80% confluence were dissociated into single cells with Accutase (Stemcell Technologies) and 8×10^5^ cells were electroporated with 2 μg Cas9-sgRNA plasmid (Addgene: #48139), 1 µg pEGFP-Puro and 200μM (6.6 μg) 180nt single-stranded DNA oligonucleotide donor template (ssODN) (PAGE-purified; Integrated DNA Technologies), using the P3 Primary Cell 4D-Nucleofector™ X Kit (Lonza), (program CA-137) on a Lonza 4D-Nucleofector™ X Unit (Lonza) according to manufacturer’s guidelines. Transfected cells were resuspended in pre-warmed Essential 8™ medium supplemented with 10 µM ROCK-inhibitor (Y-27632) (Stemcell Technologies) and seeded into two wells of a Matrigel**®** coated 6-well plate. Selection with 1 μg/ml puromycin (ThermoFisher) was commenced 24 hours post-nucleofection and continued for 24 hours. Cells were grown to confluence and passaged at low density (5×10^3^ and 1×10^4^), as single cells onto Matrigel® coated 10 cm dishes in Essential 8™ medium with 10 μM Y27632. After 8-10 days single-cell derived colonies were isolated and transferred to a Matrigel**®** coated 96-well plate. Duplicate plates were made for maintenance and restriction fragment length polymorphism (RFLP) screening. When cells for genotyping reached confluence, crude genomic DNA lysates were prepared by adding 50μl Cell Lysis Buffer (0.45% NP-40 substitute, 0.45% Tween-20, 0.2 mg/ml Proteinase K, 0.05x PCR Buffer in dH_2_O) and incubated at 55°C for 2 hours, followed by 10 minutes at 95°C. Amplicons flanking the targeting site were amplified with the following primers: Inp D447N Fw GCGGTTCTTTAGCACGGTTA and Inp D447N Rev: CTCCTCATCTCCCTCCATG using GoTaq G2 polymerase (Promega). PCR protocol: 95°C for 2 minutes; 35 cycles of 95°C for 15 seconds, 60°C for 30 seconds, 72°C for 30 seconds; and a final extension at 72°C for 5 minutes. PCR products were digested with ApoI-HF (New England Biolabs) and run on a 2% TAE agarose gel. Clones identified as carrying ApoI restriction site were evaluated for introduction of D447N mutation through Sanger sequencing (Source Bioscience). Successfully edited clones were expanded and re-sequenced and assessed for chromosomal abnormalities G-banding karyotype analysis. Quality control tests were performed after clonal passage 10 and included immunocytochemistry with a panel of antibodies to pluripotency markers (TRA-160, OCT3/4, NANOG). Standard G-banding chromosome analysis (karyotyping) was determined by TDL Genetics, London.

### Generation of cerebral organoids

Cerebral organoids were generated and maintained according to a modified Lancaster protocol (Lancaster et al., 2013) as described recently (Johnstone et al., 2019). This protocol evades the generation of embryoid bodies and goes straight to making neurospheres by dual-SMAD inhibition (Chambers et al., 2009). hiPSCs were cultured in Matrigel^®^ Matrix (Corning, 354248-10ml) coated 6-well plates in Essential 8™ Basal Medium (E8; Gibco; A15169-01) supplemented with Antibiotic-Antimycotic (100x; Invitrogen; 15240-062). The cells were grown for an average of seven days and lifted when cultures reached around 80% confluency with distinct, well defined hiPSC colonies. The colonies were lifted with a 1:1 Dispase/Collagenase enzyme mix (1mg/ml; Gibco; 17105-041 and 2 mg/ml; Gibco; 17104-019). and resuspended in 10 ml of Phase 1 medium (1:1 IMDM (Invitrogen; 41966029): Ham’s F-12 Nutrient Mix (Invitrogen; 21765029); 5g/l BSA Cohn fraction V (Europa-bioproducts; EQBAC62-0500), 1/100 Chemically Defined Lipid Concentrate (100x; Invitrogen; 11905031); 1/25000 Monothioglycerol (Sigma; M6145-100ml); 7µg/ml human Insulin (Sigma; 11376497001), 1/2000 Transferrin (Sigma; 10652202001), 1/100 Antibiotic-Antimycotic (100x; Invitrogen; 15240-062); 1mM N-acetyl cysteine (Sigma; A8199-10G); 10µM Activin Inhibitor (SB 431542; R & D systems; 1614/10) and 0.1µM LDN (Stratech; S2618)). From this point cells were cultured in suspension on an orbital shaker at 45 rpm in a cell culture CO_2_ incubator at 37°C and 5% CO_2_. After seven days the colonies were transferred to EB1 medium containing Advanced DMEM/F12 (Invitrogen; 31331028) supplemented with 1/100 Antibiotic-Antimycotic, 1/100 GlutaMAX™-I Supplement (Invitrogen; 35050-038), 1/100 N2 Supplement (Invitrogen; 17502001), 1/200 B27 Supplement (100 M; Invitrogen; 17504001) and 2.5 ng/ml Murine FGF-basic (Peprotech; 450-33). After five days, rosette forming spheres were transferred into EB2 medium for 28 days. EB2 medium consists of a 1/1 mix of Advanced DMEM/F12 and Neurobasal™ Medium (1x; Invitrogen; 21103049), supplemented with 1/100 Antibiotic-Antimycotic, 1/200 GlutaMAX™-I Supplement, 1/100 N2 Supplement, 1/200 B-27 Supplement Minus Vitamin A (50x; Invitrogen; 12587-010); 1/100 MEM Non-Essential Amino Acids Solution (10 mM; Invitrogen; 11140050) and 1.25 µg/ml human Insulin (Sigma; 11376497001). Organoids were collected for immunohistochemistry, RNA or protein extraction.

### Immunohistochemistry on organoids

For immunohistochemistry, organoids were fixed for 1 hour in 4% paraformaldehyde, incubated in 30% sucrose at +4°C for 24h, embedded in 30% sucrose/OCT mixture (1:1) and frozen on dry ice. Immunofluorescence staining was performed on 10-12 μm cryostat sections as described previously (Theil, 2005) with antibodies against mouse anti-ARL13B (Neuromab 75-287; 1:2000), rabbit anti-ARL13B (Proteintech, 1:200), mouse anti-ASCL1 (1:25, clone 24B72D11.1, BD Pharmingen #556604), rabbit anti-COUP-TFII (1:500; provided by M. Studer), rat anti-CTIP2 (1:1000, Abcam #18465), guinea pig anti-DLX2 (1:2000, Bioacademica # 74-116), rabbit anti-EMX1 (1:200; (Briata et al., 1996)), rabbit anti-FOXG1 (1:200; Abcam #18259), guinea pig anti-GLI2 (1:1000; (Cho et al., 2008)), rabbit anti-GPR161 (1:1000; 13398-1-AP), rabbit anti-GSX2 (1:200; Millipore #ABN162), rabbit anti-IFT88 (1:200; Proteintech #13967-1-AP); rabbit anti-INPP5E (1:600; Proteintech #17797-1-AP), mouse anti-ISL1/2 (1:100; DSHB clone #39.4D5), mouse anti-NKX2.1 (1:300; Abcam #ab3186), rabbit anti-OLIG2 (1:400; Millipore #AB9610), rabbit anti-PAX6 (1:400, Biolegend #901301), rabbit anti-SMO (1:600; Proteintech #20787-1-AP), rabbit anti-SOX2 (1:1000; Abcam #92494), rabbit anti-SST (1:200; Peninsula Laboratories # T-4102.0400), rabbit anti-SUFU (1:600; Proteintech #26759-1-AP), rabbit anti-TBR1 (1:400, Abcam #31940), rabbit anti-TBR2 (1:400, Abcam #23345), rabbit anti-TMEM67 (1:200; Proteintech #13975-1-AP), mouse anti-γ TUB (Sigma T6557; 1:2000), rabbit anti-TULP3 (1:600; Proteintech #13637-1-AP). Primary antibodies for immunohistochemistry were detected with Alexa- or Cy2/3-conjugated fluorescent secondary antibodies. The Tbr1 signals were amplified using biotinylated secondary IgG antibody (swine anti-rabbit IgG) (1:400, BD Biosciences) followed by Alexa Fluor 488 or 568 Streptavidin (1:100, Invitrogen). For counter staining DAPI (1:2000, Life Technologies) was used. Fluorescent and confocal images were taken on a LeicaDM 5500 B fluorescent microscope and Nikon A1R FLIM confocal microscope, respectively.

### Immunohistochemistry for pluripotency markers

hiPSCs were cultured in Matrigel^®^ Matrix (Corning, 354248-10ml) coated 24-well plates in Essential 8™ Basal Medium (E8; Gibco; A15169-01) supplemented with Antibiotic-Antimycotic (100x; Invitrogen; 15240-062). The cells were grown for an average of 4-6 days until cultures reached around 80% confluence, before they were fixed for 15 min at room temperature in 4% paraformaldehyde/DPBS. Cultures were blocked for 45 min with blocking buffer (BB) containing 6%Goat serum (Dako, S-100) in DPBS, and subsequently incubated with Mouse-anti-Tra-1-60 (1/100, Santa Cruz, sc-21705) antibody diluted in BB for one hour. Cultures were permeabilised with 0.1% Triton-X in DPBS for 10 min, followed by an overnight incubation at 4°C with Rabbit-anti-NANOG (1/800, Cell Signalling, #3580S) and Mouse-anti-OCT3/4 (1/250, Santa Cruz, sc-5279) antibodies diluted in BB supplemented with 0.1% Triton-X. Primary antibodies were detected with Goat-anti-Rabbit-488 (1/1000, Invitrogen, A11008), Goat-anti -Mouse-IgG2b-647 (1/1000, Invitrogen, A21242), Goat-anti -Mouse-IgM-555 (1/1000, Invitrogen, A21426) secondary antibodies. Fluorescent images were captured using a Zeiss observer Z1 microscope.

### In situ hybridization and qRT-PCR

In situ hybridisation on 12μm serial cryosections were performed as described previously (Theil, 2005). To generate Digoxigenin-labeled antisense probes, *GLI1* and *PTCH1* cDNAs were PCR amplified using the following oligonucleotides: 5’-TGGACTTTGATTCCCCCACCC-3’ and 5’-ATACATAGCCCCCAGCCCATAC-3’ (*GLI1*); 5’-GGTCTGCCATCCTAACACCC-3’ and 5’-CATGCTAGGTCGCCAATGGT-3’ (*PTCH1*). pBS hSHH (CT#401) was a gift from Cliff Tabin (Addgene plasmid # 13996) (Marigo et al., 1995). Images were taken on a LeicaDMLB upright compound microscope.

To validate differential expression of *PTCH1* and *GLI1*, total RNA was extracted from control and *INPP5E*^D477N/D477N^ organoids (n=3 samples per genotype) using an RNeasy Plus Micro Kit (Qiagen) and reverse transcribed using Superscript™ IV VILO™ Master ezDNase enzyme (Thermo Fisher Scientific). Quantitative reverse transcription PCR (qRT-PCR) was performed using QuantiFast SYBR Green PCR Kit (Qiagen) and a DNA Engine Opticon System (GRI); the used oligonucleotides are summarized in Supplementary Table 1. For each sample Ct values were extrapolated using the Opticon software and ratios of relative gene expression levels of *ATP5* (reference gene) and *PTCH1*/*GLI1* were calculated based on a modified ΔΔCt method taking into account different PCR kinetics (Pfaffl, 2001); PCR efficiencies are summarized in Supplementary Table 1.

### Western Blot

Protein was extracted from control and *INPP5E*^D477N/D477N^ organoids (derived from n=3 lines per genotype) as described previously (Magnani et al., 2010). 20 μg protein lysates were subjected to gel electrophoresis on a 3-8% NuPAGE® Tris-Acetate gel (Life Technologies), and protein was transferred to a Immobilon-FL membrane (Millipore), which was incubated with goat anti-h/m GLI3 (1:500, R&D Systems #AF3690) and mouse anti-β-Actin antibody (1:15000, Abcam #ab6276). After incubating with donkey anti-goat IgG IRDye680RD (1:15000, LI-COR Biosciences) and donkey anti-mouse IgG IRDye800CW secondary antibodies (1:15000, Life Technologies), signal was detected using LI-COR’s Odyssey Infrared Imaging System with Odyssey Software. Values for protein signal intensity were obtained using Image Studio Lite Version 4.0. GLI3 repressor and full-length protein levels and the GLI3 repressor/full length ratio were compared between control and mutant organoids using an unpaired t-test.

### Confocal imaging, deconvolution and image analyses

The neuroepithelia of organoids were imaged with a Nikon A1R FLIM confocal microscope with the experimenter blinded to the genotype. Laser power and gain were adjusted to maximise intensity of the staining while avoiding overexposure. The Z-stack contained between 5μm and 15 μm of tissue section imaged in 0.13 μm steps. An optical zoom of x2.26 with pixel size of 0.06 was used to show more detail of the cilia. Deconvolution was performed using Huygenes Essential with the signal to noise ratio adjusted to values between 3 and 40 and the quality threshold set to 0.01.

The fluorescence mean intensity of ciliary markers relative to axonemal ARL13B staining were analysed using ImageJ software. 15 cilia per organoid (3 organoids per genotype) were chosen that had an elongated rather than a stubby shape to prevent the accidental measurement of staining artefacts. For both, ARL13B and the marker of interest, background mean staining intensities were determined and deduced from the respective intensity levels in the cilium. The intensity ratio between the marker of interest and ARL13B was used for statistical analyses, thereby minimising bias that might have originated from a variability in the staining or image acquisition. For statistical analyses, the intensity ratios of all control and mutant organoids were collected in two separate groups. To quantify the percentage of cilia positive for a ciliary marker of interest, the number of ARL13B positive cilia that were also positive for that marker was determined using the ImageJ Cell Counting plug. 100 cilia each were counted for 3 control and 3 mutant organoids.

### Statistical Analyses

Data were analysed using GraphPadPrism 9 software with n=3-12 organoids for all analyses. Normal distribution was tested with Shapiro-Wilk or D’Agostino-Pearson omnibus normality tests and F-tests were used to test for equal variation. Normally distributed data with equal variance were analysed with unpaired t-tests but with unpaired t-tests with Welch’s correction if data showed unequal variance. In all other cases, Mann Whitney tests were used. A single asterisk indicates significance of p<0.05, two asterisks indicate significance of p<0.01, three asterisks of p<0.001 and four asterisks of p<0.0001. Graphs show the mean as well as upper and lower 95% confidence interval. Supplementary Table 2 provides a detailed summary of descriptive statistics of the tests used.

## Supporting information

Supplementary Table 2_Statistics Summary

Supplementary Table 1_Oligonucleotides

## ACKNOWLEDGEMENTS

We are grateful to Drs Thomas Becker, John Mason, Pleasantine Mill and David Price for critical comments on the manuscript. The human embryonic and fetal material was provided by the Joint MRC/Wellcome Trust (grant# MR/R006237/1) Human Developmental Biology Resource (www.hdbr.org). This work was supported by grants from a RS Macdonald Seedcord fund and from the Simons Initiative for the Developing Brain (SFARI-529085) to Thomas Theil and Siddharthan Chandran. The Chandran lab are supported by the Euan MacDonald centre for Motor Neurone Disease Research, and the UK Dementia Research Institute (DRI), which receives its funding from UK DRI Ltd., funded by the UK Medical Research Council, Alzheimer’s Society and Alzheimer’s Research UK. Bhuvaneish T. Selvaraj is a Rowling/DRI fellow, funded by the “Anne Rowling Regenerative Neurology Clinic”. Sunniva MK Bøstrand is funded by the Wellcome Trust Translational Neuroscience PhD Programme at the University of Edinburgh [108890/Z/15/Z].

## AUTHOR CONTRIBUTIONS

Conceptualisation, B.S.T., S.C., and T.T.; Investigation, L.S., A.W.,K.H.-T.,J.D.C., K.W., K.B., S.M.K.B., B.S.T, and T.T.; Writing: A.W., B.S.T., and T.T.

