## Supplementary Figure for "The ciliary gene *INPP5E* confers dorsal telencephalic identity to human cortical organoids by negatively regulating Sonic Hedgehog signalling"

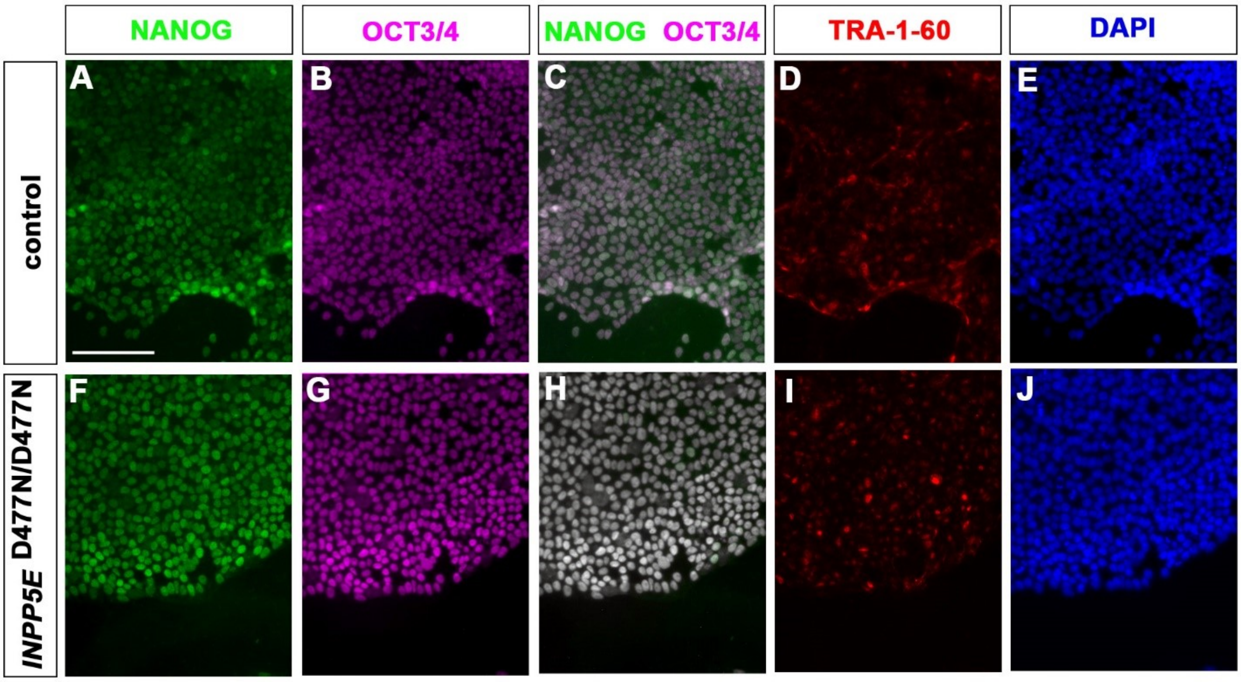


**Supplementary Figure 1: Expression of pluripotency markers in control and *INPP5E*$^{D477N/D477N}$ iPSC lines**. (A-J) Immunofluorescence stainings for the indicated markers. All iPSC lines were positive for NANOG (A, C, F, H), OCT3/4 (B, C, G, H), and TRA-1-60. Scale bar: 100 μm.
